# A Critique of Pure Learning: What Artificial Neural Networks can Learn from Animal Brains

**DOI:** 10.1101/582643

**Authors:** Anthony M. Zador

## Abstract

Over the last decade, artificial neural networks (ANNs), have undergone a revolution, catalyzed in large part by better tools for supervised learning. However, training such networks requires enormous data sets of labeled examples, whereas young animals (including humans) typically learn with few or no labeled examples. This stark contrast with biological learning has led many in the ANN community posit that instead of supervised paradigms, animals must rely instead primarily on unsupervised learning, leading the search for better unsupervised algorithms. Here we argue that much of an animal’s behavioral repertoire is not the result of clever learning algorithms—supervised or unsupervised—but arises instead from behavior programs already present at birth. These programs arise through evolution, are encoded in the genome, and emerge as a consequence of wiring up the brain. Specifically, animals are born with highly structured brain connectivity, which enables them learn very rapidly. Recognizing the importance of the highly structured connectivity suggests a path toward building ANNs capable of rapid learning.

## Introduction

Not long after the invention of computers in the 1940s, expectations were high. Many believed that computers would soon achieve or surpass human-level intelligence. Herbert Simon, a pioneer of artificial intelligence (AI), famously predicted in 1965 that “machines will be capable, within twenty years, of doing any work a man can do”—to achieve general artificial intelligence. Of course, these predictions turned out to be wildly off the mark.

In the tech world today, optimism is high again. Much of this renewed optimism stems from the impressive recent advances in artificial neural networks (ANNs) and machine learning, particularly “deep learning” ((LeCun et al., 2015)). Applications of these techniques—to machine vision, speech recognition, autonomous vehicles, machine translation and many other domains—are coming so quickly that many predict we are nearing the “technological singularity,” the moment at which artificial superintelligence triggers runaway growth and transform human civilization (Kurzweil 2005). In this scenario, as computers increase in power, it will become possible to build a machine that is more intelligent than the builders. This superintelligent machine will build an even more intelligent machine, and eventually this recursive process will accelerate until intelligence hits the limits imposed by physics or computer science.

But in spite of this progress, ANNs remain far from approaching human intelligence. ANNs can crush human opponents in games such as chess and Go, but along most dimensions—language, reasoning, common sense—they cannot approach the cognitive capabilities of a four-year old. Perhaps more striking is that ANNs remain even further from approaching the abilities of simple animals. Many of the most basic behaviors—behaviors that seem effortless to even simple animals—turn out to be deceptively challenging and out of reach for AI. In the words of one of the pioneers of AI, Hans Moravec (Moravec, 1988):

> “Encoded in the large, highly evolved sensory and motor portions of the human brain is a billion years of experience about the nature of the world and how to survive in it. The deliberate process we call reasoning is, I believe, the thinnest veneer of human thought, effective only because it is supported by this much older and much more powerful, though usually unconscious, sensorimotor knowledge. We are all prodigious Olympians in perceptual and motor areas, so good that we make the difficult look easy. Abstract thought, though, is a new trick, perhaps less than 100 thousand years old. We have not yet mastered it. It is not all that intrinsically difficult; it just seems so when we do it.”

We cannot build a machine capable of building a nest, or stalking prey, or loading a dishwasher. In many ways, AI is far from achieving the intelligence of a dog or a mouse, or even of a spider, and it does not appear that merely scaling up current approaches will achieve these goals.

The good news is that, if we do ever manage to achieve even mouse-level intelligence, human intelligence may be only a small step away. Our vertebrate ancestors, who emerged about 500 million years ago, may have had roughly the intellectual capacity of a shark. A major leap in the evolution of our intelligence was the emergence of the neocortex, the basic organization of which was already established when the first placental mammals arose about 100 million years ago (Kaas, 2011); much of human intelligence seems to derive from an elaboration of the neocortex. Modern humans (Homo sapiens) evolved only a few hundred thousand years ago—a blink in evolutionary time—suggesting that those qualities such as language and reason which we think of as uniquely human may actually be relatively easy to achieve, provided that the neural foundation is solid. Although there are genes and perhaps cell types unique to humans—just as there are for any species—there is no evidence that the human brain makes use of any fundamentally new neurobiological principles not already present in a mouse (or any other mammal), so the gap between mouse and human intelligence might be much smaller than that between than that between current AI and the mouse. This suggests that even if our eventual goal is to match (or even exceed) human intelligence, a reasonable proximal goal for AI would be to match the intelligence of a mouse.

As the name implies, ANNs were invented in an attempt to build artificial systems based on computational principles used by the nervous system (Hassabis et al., 2017). In what follows, we identify additional principles from neuroscience that might accelerate the goal of achieving artificial mouse, and eventually human, intelligence. We argue that much in contrast to ANNs, animals rely heavily on a combination of both learned and innate mechanisms. These innate processes arise through evolution, are encoded in the genome, and take the form of rules for wiring up the brain (Seung, 2012). We discuss the implications of these observations for generating next-generation machine algorithms.

### Learning by ANNs

Since the earliest days of AI, there has been a competition between two approaches: symbolic AI and ANNs. In the symbolic “good old fashion AI” approach (Haugeland, 1989) championed by Marvin Minsky and others, it is the responsibility of the programmer to explicitly program the algorithm by which the system operates. In the ANN approach, by contrast, the system “learns” from data. Symbolic AI can be seen as the psychologist’s approach—it draws inspiration from the human cognitive processing, without attempting to crack open the black box—whereas ANNs, which use neuron-like elements, take their inspiration from neuroscience. Symbolic AI was the dominant approach from the 1960s to 1980s, but since then it has been eclipsed by ANN approaches inspired by neuroscience.

Modern ANNs are very similar to their ancestors three decades ago (Rumelhart, 1986). Much of the progress can be attributed to increases in raw computer power: Simply because of Moore’s law, computers today are several orders of magnitude faster than they were a generation ago, and the application of graphics processors (GPUs) to ANNs has sped them up even more. The availability of large data sets is a second factor: Collecting the massive labeled image sets used for training would have been very challenging before the era of Google. Finally, a third reason that modern ANNs are more useful than their predecessors is that they require even less human intervention. Modern ANNs—specifically “deep networks” (LeCun et al., 2015)—learn the appropriate low-level representations (such as visual features) from data, rather than relying on hand-wiring to explicitly program them in.

In ANN research, the term “learning” has a technical usage that is different from its usage in neuroscience and psychology. In ANNs, learning refers to the process of extracting structure—statistical regularities—from input data, and encoding that structure into the parameters of the network. These network parameters contain all the information needed to specify the network. For example, a fully connected network with *N* neurons might have one parameter (e.g. a threshold) associated with each neuron, and an additional *N*^2^ parameters specifying the strengths of synaptic connections, for a total of *N* + *N*^2^ free parameters. Of course, as the number of neurons *N* becomes large, the total parameter count in a fully connected ANN is dominated by the *N*^2^ synaptic parameters.

There are three classic paradigms for extracting structure from data, and encoding that structure into network parameters (i.e. weights and thresholds). In *supervised* learning, the data consist of pairs—an input item (e.g. an image) and its label (e.g. the word “giraffe”)—and the goal is to find network parameters that generate the correct label for novel pairs. In *unsupervised* learning, the data have no labels; the goal is to discover statistical regularities in the data without explicit guidance about what kind of regularities to look for. For example, one could imagine that with enough examples of giraffes and elephants, one might eventually infer the existence of two classes of animals, without the need to have them explicitly labeled. Finally, in *reinforcement* learning, data are used to drive actions, and the success of those actions is evaluated based on a “reward” signal.

Much of the progress in ANNs has been in developing better tools for supervised learning. A central consideration in supervised learning is “generalization.” As the number of parameters increases, so too does that “expressive power” of the network—the complexity of the input-output mappings that the network can handle. A network with enough free parameters can fit any function (Cybenko, 1989; Hornik, 1991), but the amount of data required to train a network without overfitting generally also scales with the number of parameters. If a network has too many free parameters, the network risks “overfitting” data, i.e. it will generate the correct responses on the training set of labeled examples, but will fail to generalize to novel examples. In ANN research, this tension between the flexibility of a network (which scales with the number of neurons and connections) and the amount of data needed to train the network (more neurons and connections generally require more data) is called the “bias-variance tradeoff” (Figure 1). A network with more flexibility is more powerful, but without sufficient training data the predictions that network makes on novel test examples might be wildly incorrect—far worse than the predictions of a simpler, less powerful network. To paraphrase “Spiderman”: With great power comes great responsibility (to obtain enough labeled training data) (Lee, 1962). The bias-variance tradeoff explains why large networks require large amounts of labeled training data.

**Figure 1:**
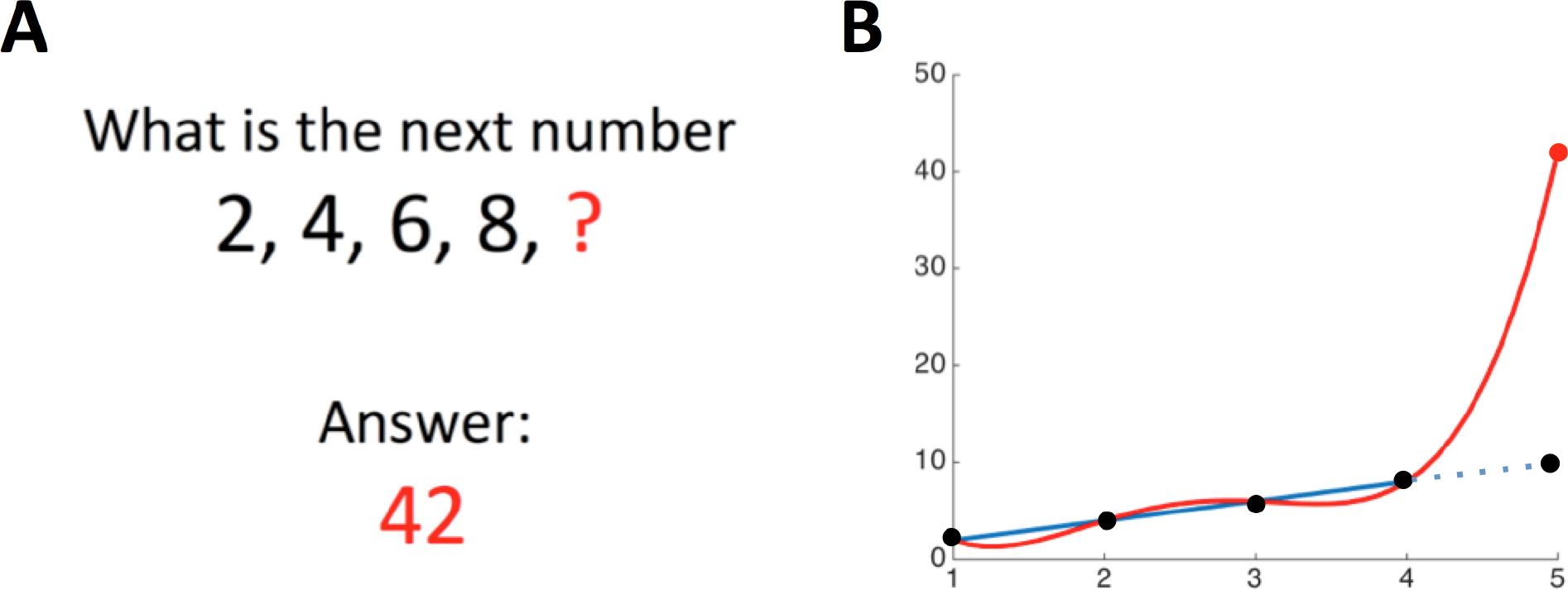
The “bias-variance tradeoff” in machine learning can be seen as a formalization of Occam’s Razor. (A) As an example of the bias-variance tradeoff and the risks of overfitting, consider the following puzzle: Find the next point in the sequence {2, 4, 6, 8, ?}. Although the natural answer may seem to be 10, a fitting function consisting of polynomials of degree 4—a function with 5 free parameters—might very well predict that the answer is 42. (B) The reason is that in general, it takes two points to fit a line, and *k* + 1 points to fit the coefficients *c_i_* of a polynomial *f* (*x*) = *c*_0_ + *c*_1_*x* + … + *c_k_x^k^* of degree *k*. Since we only have data for 4 points, the next entry could be literally any number (e.g. red curve). To get the expected answer, 10, we might restrict the fitting functions to something simpler, like lines, by discouraging the inclusion of higher order terms in the polynomial (blue line).

### Learning by animals

The term “learning” in neuroscience (and in psychology) refers to a long-lasting change in behavior that is the result of experience. Learning in this context encompasses animal paradigms such as classical and operant conditioning, as well as an array of other paradigms such as learning by observation or by instruction. Although there is some overlap between the neuroscience and ANN usage of learning, in some cases the terms differ enough to lead to confusion.

Perhaps the greatest divergence in usage is the application of the term “supervised learning.” Supervised learning is central to the many successful recent applications of ANNs to real-world problems of interest. For example, supervised learning is the paradigm that allows ANNs to categorize images accurately. However, to ensure generalization, training such networks requires enormous data sets; one visual query system was trained on more than 10^7^ “labeled” examples (question-answer pairs) (Antol et al., 2015). Although the final result of this training is an ANN with a capability that, superficially at least, mimics the human ability to categorize images, the process by which the artificial system learns bears little resemblance to that by which a newborn learns. There are only 10^7^ seconds in a year, so a child would need to ask one question every second of her life to receive a comparable volume of labeled data; and of course, most images encountered by a child are not labeled. There is, thus, a mismatch between the available pool of labeled data and how quickly children learn. Clearly, children do not rely mainly on supervised algorithms to learn to categorize objects.

Considerations such as these have motivated the search in the machine learning community for more powerful learning algorithms, for the “secret sauce” posited to enable children to learn how to navigate the world within a few years. Many in the ANN community posit that instead of supervised paradigms, we rely instead primarily on unsupervised paradigms to construct representations of the world. Because unsupervised algorithms do not require labeled data, they could potentially exploit the tremendous amount of raw (unlabeled) sensory data we receive. Indeed, there are several unsupervised algorithms which generate representations reminiscent of those found in the visual system (Bell and Sejnowski, 1997; Olshausen and Field, 1996; van Hateren and Ruderman, 1998). Although at present these unsupervised algorithms are not able to generate visual representations as efficiently as supervised algorithms, there is no known theoretical principle or bound that precludes the existence of such an algorithm (although the No-Free-Lunch theorem for learning algorithms (Wolpert, 1996) states that no completely general-purpose learning algorithm can exist, in the sense that for every learning model there is a data distribution on which it will fare poorly). Every learning model must contain implicit or explicit restrictions on the class of functions that it can learn. Thus, while the number of labeled images a child encounters during her first 10^7^ seconds of life might be small, the total sensory input received during that time is quite large; perhaps Nature has evolved a powerful unsupervised algorithm to exploit this large pool of data. Discovering such an unsupervised algorithm—if it exists—would lay the foundation for a next generation of ANNs.

### Learned and innate behavior in animals

A central question, then, is how animals function so well so soon after birth, without the benefit of massive supervised training data sets. It is conceivable that unsupervised learning, exploiting algorithms more powerful than any yet discovered, may play a role establishing sensory representations and driving behavior. But even such a hypothetical unsupervised learning algorithm is unlikely to be the whole story. Indeed, the challenge faced by this hypothetical algorithm is even greater than it appears. Humans are an outlier: We spend more time learning than perhaps any other animal, in the sense that we have an extended period of immaturity. Many animals function effectively after 10^6^, 10^5^, or even fewer seconds of life: A squirrel can jump from tree to tree within months of birth, a colt can walk within hours, and spiders are born ready to hunt. Examples like these suggest that the challenge may exceed the capacities of even the cleverest unsupervised algorithms.

So if unsupervised mechanisms alone cannot explain how animals function so effectively at (or soon after) birth, what is the alternative? The answer is that much of our sensory representations and behavior are largely innate. For example, many olfactory stimuli are innately attractive or appetitive (blood for a shark (Yopak et al., 2015) or aversive (fox urine for a rat (Apfelbach et al., 2005)). Responses to visual stimuli can also be innate. For example, mice respond defensively to looming stimuli, which may allow for the rapid detection and avoidance of aerial predators (Yilmaz and Meister, 2013). But the role of innate mechanisms goes beyond simply establishing responses to sensory representations. Indeed, most of the behavioral repertoire of insects and other short-lived animals is innate. There are also many examples of complex innate behaviors in vertebrates, for example in courtship rituals (Tinbergen, 1951). A striking example of a complex innate behavior in mammals is burrowing: Closely related species of deer mice differ dramatically in the burrows they build with respect to the length and complexity of the tunnels (Weber and Hoekstra, 2009; Metz et al., 2017). These innate tendencies are independent of parenting: Mice of one species reared by foster mothers of the other species build burrows like those of their biological parents. Thus, it appears that a large component of an animal’s behavioral repertoire is not the result of clever learning algorithms—supervised or unsupervised—but rather of behavior programs already present at birth.

From an evolutionary point of view, it is clear why innate behaviors are advantageous. The survival of an animal requires that it solve the so-called “four Fs”—feeding, fighting, fleeing, and mating—repeatedly, with perhaps only minor tweaks. Each individual is born, and has a very limited time—from a few days to a few years—to figure out how to solve these four problems. If it succeeds, it passes along part of its solution (i.e. half its genome) to the next generation. Consider a species X that achieves at 98% of its mature performance at birth, and its competitor Y that functions only at 50% at birth, requiring a month of learning to achieve mature performance. (Performance here is taken as some measure of fitness, i.e. ability of an individual to survive and propagate). All other things being equal (e.g., assuming that mature performance level is the same for the two species), species X will outcompete species Y, because of shorter generation times and because a larger fraction of individuals survive the first month to reproduce (Fig. 2A).

**Figure 2:**
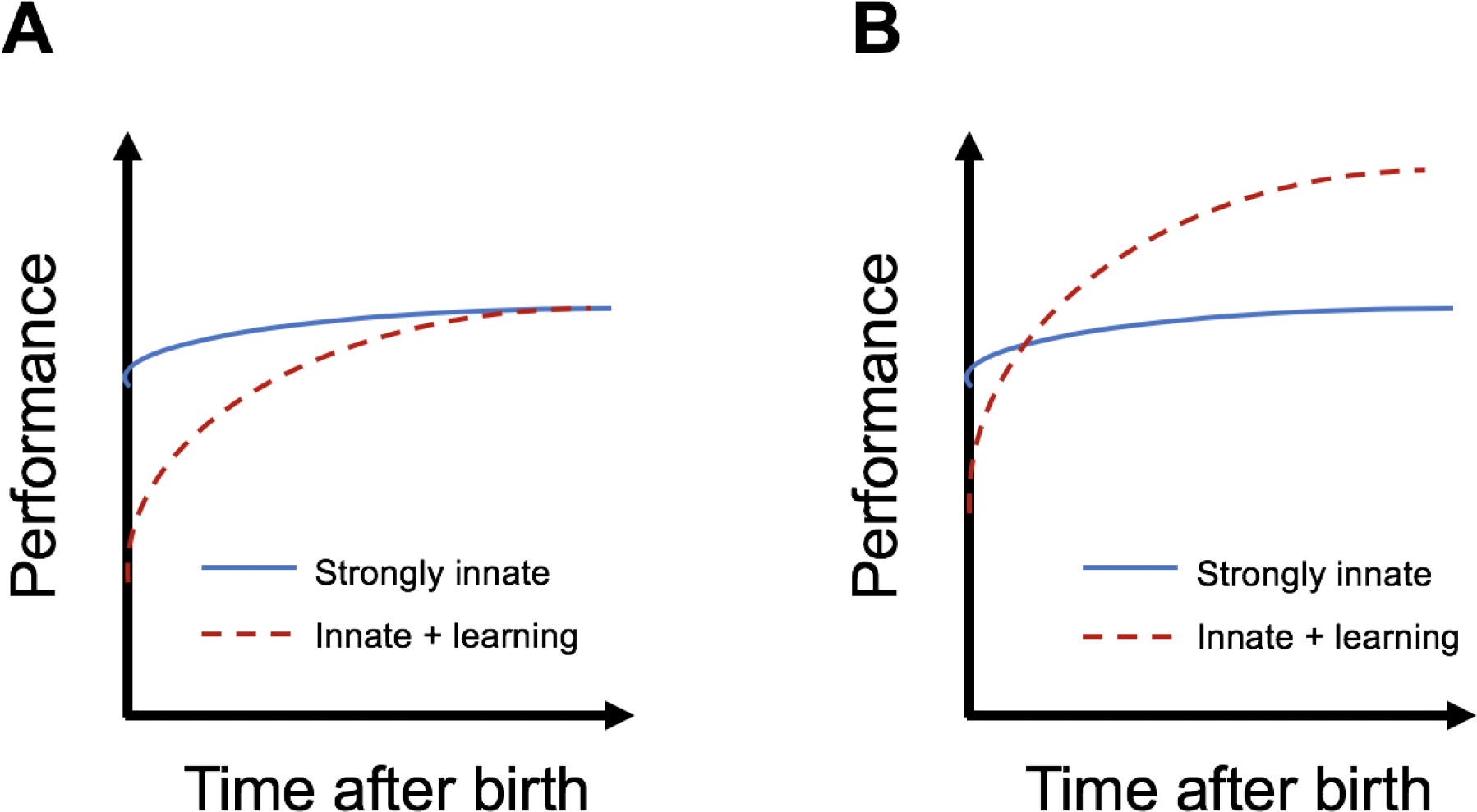
Evolutionary tradeoff between innate and learning strategies. (A) Two species differ in their reliance on learning, and achieve the same level of fitness. All other things being equal, the species relying on a strongly innate strategy will outcompete the species employing a mixed strategy. (B) A species using the mixed strategy may thrive if that strategy achieves a higher asymptotic level of performance.

In general, however, all other things may not be equal. The mature performance achievable via purely innate mechanisms might not be the same as that achievable with additional learning (Fig. 2A). If an environment is changing rapidly—e.g. on the timescale of a single individual—innate behavioral strategies might not provide a path to as high a level of mature performance as a mixed strategy that relies in part on learning. For example, a fruit-eating animal might evolve an innate tendency to look for trees; but the locations of the fruit groves in its specific environment must be learned by each individual. There is, thus, pressure to evolve an appropriate tradeoff between innate and learned behavioral strategies, reminiscent of the bias-variance tradeoff in supervised learning.

### Innate and learned behaviors are synergistic

The line between innate and learned behaviors is, of course, not sharp. Innate and learned behaviors and representations interact, often synergistically. For example, rodents and other animals form a representation of space—a “cognitive map”—in the hippocampus. This representation consists of place cells, which become active when the animal enters a particular place in its environment known as a “place field.” A given place cell typically has only one (or a few) place fields in a particular environment. The propensity to form place fields is innate: A map of space emerges when young rat pups explore an open environment outside the nest for the very first time (Langston et al., 2010). However, the content of place fields is learned; indeed, it is highly labile, since new place fields form whenever the animal enters a new environment. Thus, the scaffolding for the cognitive map is innate, but the specific maps constructed on this scaffolding are learned.

This form of synergy between innateness and learning is common. For example, human infants can discriminate faces soon after birth, and monkeys raised with no exposure to faces show a preference for faces upon first exposure, reflecting the contribution of innate mechanisms to face salience and perception (McKone et al., 2009). In human and non-human primates there exists a specific cortical area, the FFA (fusiform face area), which is selectively engaged in the perception of faces; patients with focal loss of the FFA suffer a permanent deficit in face processing (Kanwisher and Yovel, 2006). However, the specific faces recognized by each individual are learned during the course of that individual’s lifetime. Thus, as with place cells in the hippocampus, the innate circuitry for processing faces may provide the scaffolding, but the specific faces that populate this scaffolding are learned. Similar synergy may accelerate the acquisition of language by children: The innate circuitry in areas like Wernicke’s and Broca’s may provide the scaffolding, enabling the specific syntax and vocabulary of any specific language to be learned rapidly (Pinker, 1994; Marcus, 2004). This synergy between innate and learned behavior could arise from evolutionary pressure of the sort depicted in Figure 2B.

### Genomes specify rules for brain wiring

We have argued that the main reason that animals function so well so soon after birth is that they rely heavily on innate mechanisms. Innate mechanisms, rather than heretofore undiscovered unsupervised learning algorithms, provide the base for Nature’s secret sauce. These innate mechanisms are encoded in the genome. Specifically, the genome encodes blueprints for wiring up their nervous system, where by wiring we refer to both the specification of which neurons are connected, as well as to the strengths of those connections. These blueprints have been selected by evolution over hundreds of millions of years, operating on countless quadrillions of individuals. The circuits specified by these blueprints provide the scaffolding for innate behaviors, as well as for any learning that occurs during an animal’s lifetime.

If the secret sauce is in our genomes, then we must ask what exactly our genomes specify about wiring. In some simple organisms, genomes specify every connection between every neuron, to the minutest detail. The simple worm c. elegans, for example, has 302 neurons and about 7000 synapses; in each individual of an inbred strain, the wiring pattern is exactly the same (Chen et al., 2006). So, at one extreme, the genome can encode a lookup table, which is then transformed by developmental processes into a circuit with precise and stereotyped connections.

But in larger brains, such as those of mammals, synaptic connections cannot be specified so precisely; the genome simply does not have sufficient capacity to specify every connection explicitly. The human genome has about 3 × 10^9^ nucleotides, so it can encode no more than about 1 GB of information—an hour or so of streaming video (Wei et al., 2013). But the human brain has about 10^11^ neurons, and more than 10^3^ synapses per neuron. Since specifying a connection target requires about log_2_ 10^11^ = 35 bits/synapse, it would take about 3.5 × 10^15^ bits to specify all 10^14^ connections. (This may represent an underestimate because it considers only the presence or absence of a connection; a few extra bits/synapse would be required to specify graded synaptic strengths. But because of synaptic noise and for other reasons, synaptic strength may not be specified very precisely. So, in large and sparsely connected brains, most of the information is probably needed to specify the locations the nonzero elements of the connection matrix rather than their precise value.). Thus, even if every nucleotide of the human genome were devoted to efficiently specifying brain connections, the information capacity would still be at least six orders of magnitude too small.

These fundamental considerations explain why in most brains, the genome cannot specify the explicit wiring diagram, but must instead specify a set of rules for wiring up the brain during development. Even a short set of rules can readily specify the wiring of a very large number of neurons; in the limit, a nervous system wired up like a grid would require only the single rule that each neuron connect to its 4 nearest neighbors (although such a nervous system would probably not be very interesting). Another simple rule, but which yields a much more interesting result, is: Given the rules of the game, find the network that plays Go as well as possible (Silver et al., 2018). In practice, the circuits found in animal brains often seem to rely on repeating modules. There has long been speculation that the neocortex consists of many copies of a basic “canonical” microcircuit (Douglas et al., 1989; Harris and Shepherd, 2015), which are wired together to form the entire cortex.

### Supervised learning or supervised evolution?

As noted above, the term “learning” is used differently in ANNs and neuroscience. At the most abstract level, learning can be defined as the process of encoding statistical regularities from the outside world into the parameters (mostly connection weights) of the network. But in the context of animal learning, the source of the input data for learning is limited only to the animal’s “experience,” i.e. to those events that occur during the animal’s lifetime. Wiring rules encoded in the genome that do not depend on experience, such as those used to wire up the retina, are not usually termed “learning.” Because the terms “lifetime” and “experience” are not well defined when applied to an ANN, reconciling the two definitions of learning in ANNs vs. neuroscience poses a challenge.

If, as we have argued above, much of an animal’s behavior is innate, then an animal’s life experiences represent only a small fraction of the data that contribute to its fitness; another potentially much larger pool of data contributes to its innate behaviors and representations. These innate behaviors and representations arise through evolution by natural selection. Thus evolution, like learning, can also be viewed as a mechanism for extracting statistical regularities, albeit on a much longer time scale than learning. Evolution can be thought of as a kind of reinforcement algorithm, operating on the timescale of generations, where the reinforcement signal consists of the number of progeny an individual generates. ANNs are engaged in an optimization process that must mimic both what is learned during evolution and the process of learning within a lifetime, whereas for animals learning only refers to within lifetime changes.

In this view, supervised learning in ANNs should not be viewed as the analog of learning in animals. Instead, since most of the data that contribute an animal’s fitness are encoded by evolution into the genome, it would perhaps be just as accurate (or inaccurate) to call rename it “supervised evolution.” Such a renaming would emphasize that “supervised learning” in ANNs is really recapitulating the extraction of statistical regularities that occurs in animals by both evolution and learning. In animals, there are two nested optimization processes: an outer “evolution” loop acting on a generational timescale, and an inner “learning” loop, which acts on the lifetime of a single. Supervised (artificial) evolution may be much faster than natural evolution, which succeeds only because it can benefit from the enormous amount of data represented by the life experiences of quadrillions of individuals over hundreds of millions of years.

Although there are parallels between learning and evolution, there are also important differences. Notably, whereas learning can act directly on synaptic weights through Hebbian and other mechanisms, evolution acts on brain wiring only indirectly, through the genome. The genome doesn’t encode representations or behaviors directly; it encodes wiring rules and connection motifs. The limited capacity of the mammalian genome—orders of magnitude smaller than would be needed to specify all connections explicitly—may act as a regularizer (Poggio et al., 1985) or an information bottleneck (Tishby et al., 2000), shifting the balance from variance to bias. In this regard, it is interesting to note that the size of the genome itself is not a fixed constraint, but is itself highly variable across species. The size of the human genome is about average for mammals, but dwarfed in size by that of many fish and amphibians; the lungfish of the marbled genome is more than 40 times larger than that the humans (Leitch, 2007). The fact that the human genome could potentially have been much larger suggests that the regularizing effect imposed by the limited capacity of the genome might represent a feature rather than a bug.

### Implications for ANNs

We have argued that animals are able to function well so soon after birth because they are born with highly structured brain connectivity. This connectivity provides a scaffolding upon which rapid learning can occur. Innate mechanisms thus work synergistically with learning. We suggest that analogous approaches might inspire new approaches to accelerate progress in ANNs.

The first lesson from neuroscience is that much of animal behavior is innate, and does not arise from learning. Animal brains are not the blank slates, equipped with a general purpose learning algorithm ready to learn anything, as envisioned by some AI researchers; there is strong selection pressure for animals to restrict their learning to just what is needed for their survival (Fig. 2). The idea that animals are predisposed to learn certain things rapidly is related to the idea of “meta-learning” in AI research (Andrychowicz et al., 2016; Finn et al., 2017; Bellec et al., 2018), and related ideas in cognitive science (Tenenbaum et al., 2011). In this formulation, there is an outer loop (e.g. evolution) which optimizes learning mechanisms to have inductive biases that allow us to learn very specific things very quickly.

The importance of innate mechanisms suggests that an ANN solving a new problem should attempt as much as possible to build on the solutions to previous related problems. Indeed, this idea is related to an active area of research in ANNs, “transfer learning,” in which connections pre-trained in the solution to one task are transferred to accelerate learning on a related task (Pan and Yang, 2010; Vanschoren, 2018). For example, a network trained to classify objects such as elephants and giraffes might be used as a starting point for a network that distinguishes trees or cars. However, transfer learning differs from the innate mechanisms used in brains in an important way. Whereas in transfer learning the ANN’s entire connection matrix (or a significant fraction of it) is typically used as a starting point, in animal brains the amount of information “transferred” from generation to generation is smaller, because it must pass through the bottleneck of the genome. Passing the information through the genomic bottleneck may select for wiring rules that are more generic. For example, the wiring of the visual cortex is quite similar to that of the auditory cortex (although each area has idiosyncrasies (Oviedo et al., 2010). This suggests that the hypothesized canonical cortical circuit provides, with perhaps only minor variations, a foundation for the wide variety of tasks that mammals perform. Neuroscience suggests that there may exist more powerful mechanisms—a kind of generalization of transfer learning—which operate not only within a single sensory modality like vision, but across sensory modalities and even beyond.

A second lesson from neuroscience follows from the fact that the genome doesn’t encode representations or behaviors directly or optimization principles directly. The genome encodes wiring rules and patterns, which then must instantiate behaviors and representations. It is these wiring rules that are the target of evolution. This suggests wiring topology and network architecture as a target for optimization in artificial systems. Classical ANNs largely ignored the details of network architecture, guided perhaps by theoretical results on the universality of fully connected three-layer networks (Cybenko, 1989; Hornik, 1991). But of course, one of the major advances in the modern era of ANNs has been convolutional neural networks (CNNs), which use highly constrained wiring to exploit the fact that the visual world is translation invariant (LeCun et al., 1989, 1998). The inspiration for this revolutionary technology was in part the structure of visual receptive fields (Fukushima, 1980). This is the kind of innate constraint that in animals would be expected to arise through evolution (Real et al., 2017); there might be many others yet to be discovered. Other constraints on wiring and learning rules are sometimes imposed in ANNs through hyperparameters, and there is an extensive literature on hyperparameter optimization. At present, however, ANNs exploit only a tiny fraction of possible network architectures, raising the possibility that more powerful, cortically-inspired architectures remain to be discovered.

In principle, wiring motifs could be discovered by experimental neuroscience, through analysis of brain connectivity. Unfortunately, we still have only a very incomplete understanding of the details of cortical wiring, because we have lacked the experimental tools for studying neuronal wiring at sufficient throughput and resolution. However, significant recent advances in experimental methods may soon provide the necessary details. Local circuitry can be determined with serial electron microscopy; there is now an ambitious project to determine every synapse within a 1 mm^3^ cube of mouse visual cortex (Strickland, 2017). Long-range projections can be determined in a high-throughput manner using MAPseq (Kebschull et al., 2016) or by other methods. Thus the details of cortical wiring may soon be available, and provide an experimental basis for ANNs.

## Conclusions

The notion that the brain provides insights for AI is not novel; indeed, it is at the very foundation of ANN research. ANNs represented an attempt to capture some key aspects of the nervous system: many simple units, connected by synapses, operating in parallel. Several subsequent advances also arose from neuroscience. For example, the reinforcement learning algorithms underlying recent successes such as AlphaGo Zero (Silver et al., 2018) draw their inspiration from the study of animal learning. Similarly, CNNs were inspired by the structure of the visual cortex.

But it remains controversial whether further progress in AI will benefit from the study of animal brains. Perhaps we have learned all that we need to from animal brains. Just as airplanes are very different from birds, so one could imagine that an intelligent machine would operate by very different principles from those of a biological organism. We argue that this is unlikely because what we demand from an intelligent machine—what is sometimes misleadingly called “artificial general intelligence”—is not general at all; it is highly constrained to match human capacities so tightly that only a machine structured similarly to a brain can achieve it. An airplane is by some measures vastly superior to a bird: It can fly much faster, at greater altitude, for longer distances, with vastly greater capacity for cargo. But a plane cannot dive into the water to catch a fish, or swoop silently from a tree to catch a mouse. In the same way, modern computers have already by some measures vastly exceeded human computational abilities (e.g. chess), but cannot match humans on the decidedly specialized set of tasks defined as general intelligence. If we want to design a system that can do what we do, we will need to build it according to the same design principles.

## Acknowledgements

We thank Adam Kepecs, Alex Koulakov, Alex Vaughan, Barak Pearlmutter, Blake Richards, Dan Ruderman, Daphne Bavelier, David Markowitz, Konrad Koerding, Mike Deweese, Peter Dayan, Petr Znamenskiy, and Ryan Fong for many thoughtful comments on and discussions about earlier versions of this manuscript.

